# A user-friendly input–output (IO) curve analysis tool for variable brain stimulation responses, particularly evoked potentials in transcranial magnetic stimulation (TMS)

**DOI:** 10.1101/2024.01.12.575452

**Authors:** Ke Ma, Stephan M. Goetz

## Abstract

**Background:** Motor-evoked potentials (MEPs) are among the few readily observable direct responses to suprathreshold stimuli administered to the brain. They serve for a variety of applications, often in the form of dose–response curves, also called recruitment or input–output (IO) curves. However, MEPs and thus IO curves demonstrate extreme trial-to-trial variability that can exceed more than two decimal orders of magnitude. Recent studies have identified issues in previous statistical analysis of IO curves and introduced better methods, others could quantitatively separate several widely independent variability sources. However, research has failed providing the field with a user-friendly implementation of the methods for analysing such IO curves statistically sound and separating variability so that they were limited to a few research groups.

**Objective:** This work intends to provide the latest methods for analysing IO curves and extract variability information in an open-source package so that the community can easily use and adapt them to their own needs.

**Methods:** We implemented recent IO curve methods with a graphical user interface and provided the code as well as compiled versions for Mac, Linux, and Windows.

**Results:** The application imports typical IO data of individual stimulus–response sets, guides users step by step through the analysis, and allows exporting of the results including figures for post-hoc analysis and documentation.

**Conclusion:** This graphical application offers a user-friendly environment for analysing the variability of evoked potentials and its various contributions, catering to the needs of clinical and experimental neuroscientists.

## Introduction

Motor-evoked potentials (MEPs) are among the few directly observable acute responses to brain stimulation, e.g., transcranial magnetic stimulation (TMS) of the motor cortex [1]. They therefore serve for many applications, such as for probing excitability changes through neuromodulation or as a reference through the motor threshold for almost all other stimulation procedures. The motor input–output (IO) curve is a well-known and important detection measurement for assessing changes in the motor system [2]. However, the MEP following brain and certain types of spinal stimulation can exhibit high levels of intra-individual trial-to-trial variability and shows rather intricate, highly skewed, heteroscedastic distributions [3, 4, 5, 6, 7, 8]. Therefore, simple least-square fitting of sigmoidal curves, which lead to asymmetric residuals and are prone to spurious results, cause systematic errors and risk spurious biomarkers. For a notably better, though still not perfect consideration of the skewed distributions, normalisation of the distribution was strongly recommended [9, 10]. Logarithmic normalisation improves stability and reduces model bias.

Beyond normalisation, more sophisticated models were able to reproduce all previously not explicable distribution features. They could further identify and isolate several apparently widely independent sources of variability affecting IO curves, one with strong stimulus-strength dependence affecting the curve in the x direction in IO curve plots and the other rather independent of the stimulation and multiplicatively changing the MEPs in y direction [11]. These models can quantitatively derive the size of both variability sources, which represent and combine different mechanisms. Coil position fluctuations, for instance, would be exclusively represented by the first variability source, i.e., the one affecting the graphical curve in x direction. Later research refined the model class and confirmed its statistical better description of the observed behaviour, including phenomena such as changing distribution spread and skewness along the stimulus strength in contrast to the conventional least-square regression model, in a larger dataset [12]. This model class appears to apply to any form of stimulation of neural circuits, including electrical stimulation [13]. Furthermore, such models applied to several stimulation methods demonstrated that coil position fluctuations are not the only and—in case of well controlled experimental methods—not even dominant source of variability subsumed in the first variability term.

This statistical model class could assist us in analytically extracting hidden information of variability from the MEP measurements, quantitatively describing the variability, and technically reducing the unknown variability. Although previous methods are well documented in the literature and commented codes are shared in the community, they may not be helpful for many users, who would rather appreciate a readily usable computer programme. Furthermore, such a programme should ideally be available in code for critical review and to enable improvements as well as amendments by other researchers. Therefore, we implemented and combined the available methods in several MATLAB classes (R2022b, The Mathworks, USA) following an object-oriented programming style and further developed a graphical user interface.

## Programme and User Interface

The IO curve analyser guides users through the basis functionalities. Figure 1 **(C)** outlines the entire application control flow. Potential users can easily follow the flowchart to use this application, and additionally, this application also includes an intuitive step-by-step tutorial program.

**Figure 1:**
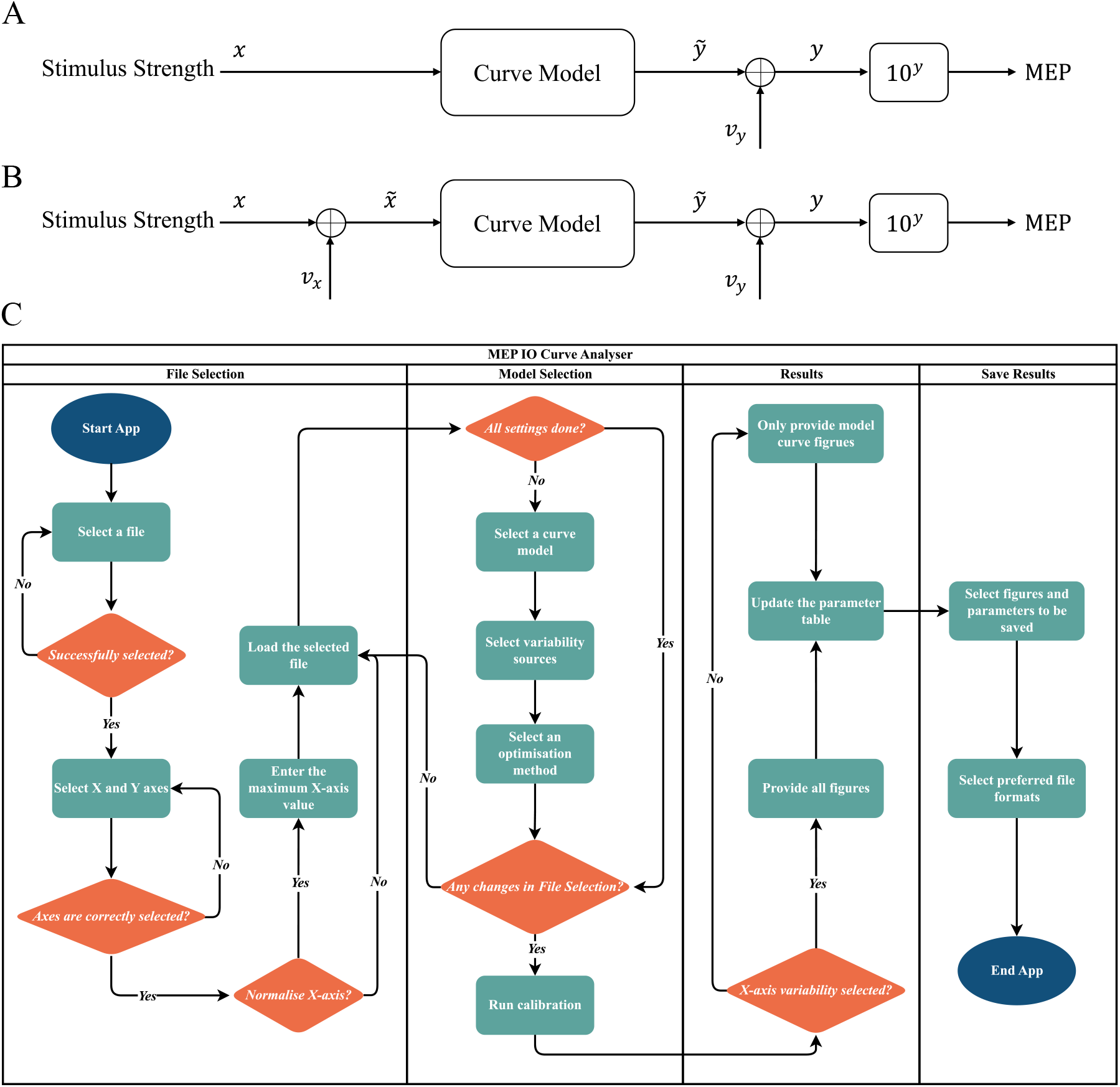
Fundamental structure of the models and the overall IO-curve analyser application. **(A)** The simpler implemented model considers only one variability source at the output and the exponential quality of the curve through log-normalisation. **(B)** Following recent developments, the advanced model adds a further variability source at the input, which can quantify excitability and coil position fluctuations even on the individual stimulus level. **(C)** The overall application has four functional components: *File Selection, Model Selection, Results*, and *Save Results*. The cyan boxes represents the processing, the orange diamonds require decisions, and dark blue boxes stand for Start/End. The output *y* is transformed using an exponential function with base 10 in both **(A)** and **(B)**.

Initially, users select their dataset file, which can be two-column comma-separated value files or Excel spreadsheets with one column representing the stimulation strengths of the pulses and the other one the corresponding responses, typically as peak-to-peak MEP values, e.g., in volts. The user can select the MEP amplitude variable from the various identified columns in the file (y axis) and the corresponding stimulus strength (x axis). The application automatically checks if the same axis is repeatedly selected. Given that the units and ranges of stimulus strength (e.g., 0 to 1, 0 to 100 percent maximum machine output, millivolts, or A/μs current slopes in TMS; certain milliampere values for electrical stimulation) can vary among different brain/spinal stimulation experiments, we provide an option of normalisation on the x axis by defining a maximum x axis value, which will become the maximum value after normalisation.

After the file is successfully loaded, users can select a desired curve model for describing the MEP IO relationship. All models use log-transformation to the base of 10 for the MEP amplitude [10]. At present, a Hill-type function with five parameters

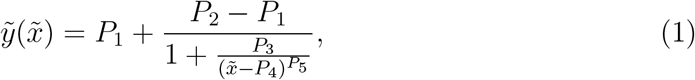

where 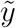 is the log-transformed response, and **P** = [*P*_1_, *P*_2_, *P*_3_, *P*_4_, *P*_5_] is the shape parameter vector (see details in Goetz and Peterchev [11]) serving as the standard sigmoid. Per default, the statistical model considers only one variability along the y direction, described by its standard deviation *v*_y_, which represents the most widely used nonlinear curve regression shown in Figure 1 **(A)** [9, 10, 14, 15]. This model represents the more widely used IO curve regression and analysis method that accounts for the log-normal distribution of the MEP variability [9, 10, 14, 15].

Users can also add an additional variability source (*v*_x_), which affects the effective stimulus strength and therefore varies the IO curve along the x axis in the IO graph (Figure 1 **(B)**). Both sources individually are Gaussian and independent in the model [11]. If only one variability term is selected, the regression turns into a more conventional nonlinear curve regression. Both cases, i.e., a single variability source or allowing two variability sources, perform a maximum-likelihood estimate of the best matching sigmoid parameters and variability sizes. Thus, the methods also quantify the trial-to-trial variability, which could serve as biomarkers, e.g., for pharmaceuticals.

The *Model Selection* frame of the user interface provides several commonly used optimisation methods. The global particle swarm method is very robust with respect to the initial parameter guesses but can be computationally intensive. The alternative simplex and interior point methods are convex optimisation methods, which converge substantially faster but may get stuck in local minima and can be very sensitive to the selection of the initial parameters. The programme runs in any case a standard least-square regression for the selected sigmoid curve to set the initial point of the shape parameter vector for the subsequent maximum-likelihood estimation with either one or two variability sources. The user can furthermore control the maximum number of iterations and function evaluations, as well as the target tolerances for the likelihood and the parameters. Those parameters determine the duration of the regression and the numerical accuracy of the result.

After the regression, the IO curve analyser updates the identified curve and variability parameters in the table and plots the stimulus–response pairs, curves, and tolerance ranges in the *Results* frame of the user interface. There are three representative figures. The first is an MEP IO curve with variability ranges calibrated using maximum-likelihood estimation (see details in Goetz and Peterchev [11]). We also offer an excitation level plot to indicate if the stimulated target was more or less excitable compared to their temporal average and the corresponding p-values as a plot to provide some statistical handle on the level of an individual stimulus (see Goetz et al. [13] for details). The plots of the excitation level and its corresponding p-value are not available when *v*_x_ is not selected so that only one variability source is considered, as they depend on the estimation of *v*_x_. The results and plots can be stored in various formats for further analysis and documentation.

The object-oriented design of the programme allows a relatively simple expansion, e.g., with own models. The authors invite their colleagues to make their modifications and additions available to the community to improve the available methodology and tools.

## Results and Discussion

Figure 2 presents the layout of the main window of this application, including its three frames of input file selection, model details, and results. The save button opens a pop-up window with a wide variety of options. The *File Selection* frame allows users to select data files and choose x-axis and y-axis. In the *Model Selection* frame, available curve models and optimisation methods are separately integrated into two drop-down lists. Additionally, the mathematical equation of the chosen curve model will be displayed in the text box below. Moreover, the *Results* section presents the calibrated parameters for the MEP IO curve using maximum-likelihood estimation and shows the aforementioned three plots.

**Figure 2:**
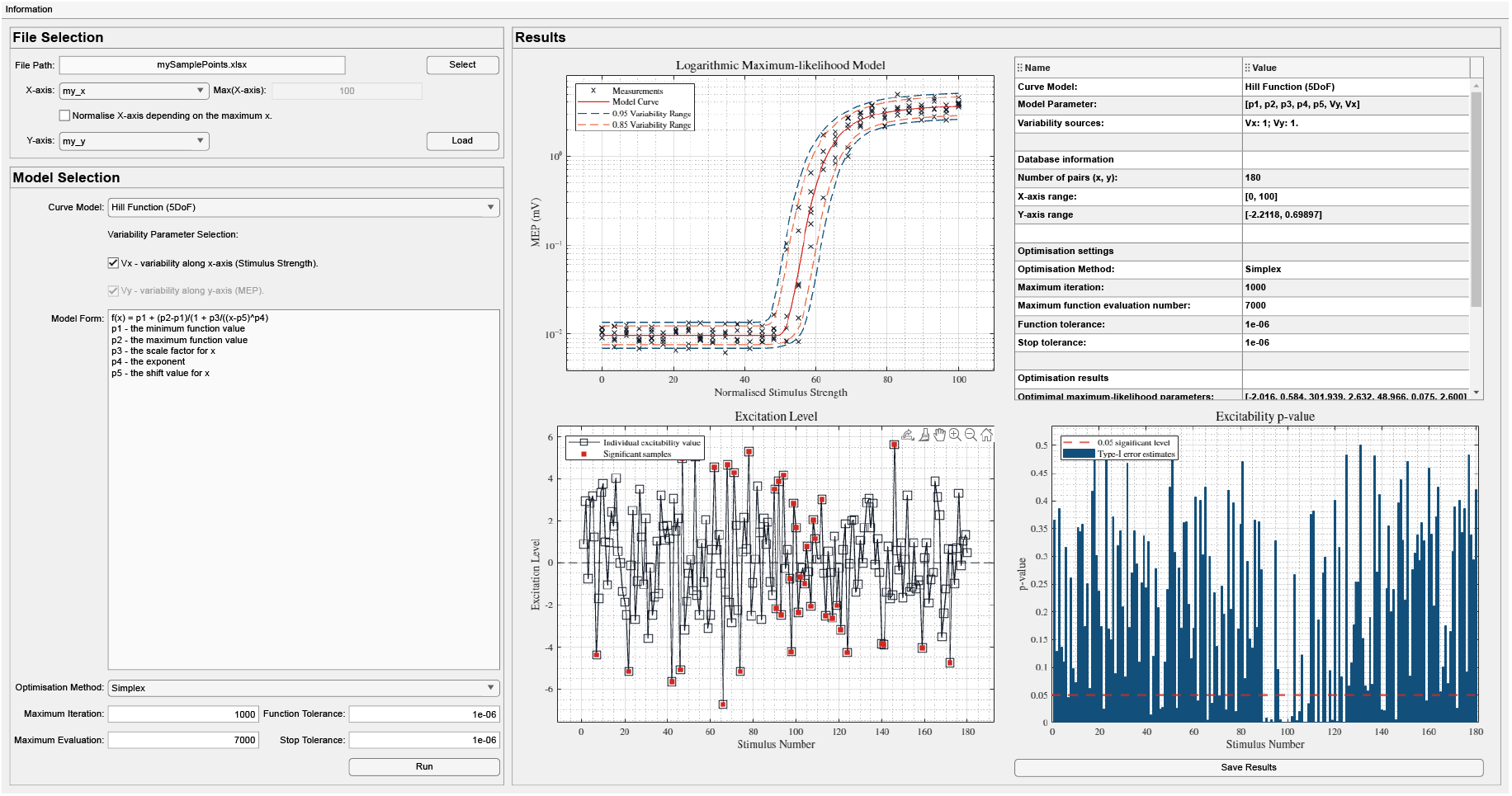
The layout of the *MEP IO Curve Analyser* application. The main window structure reflects with its frames the functionality from Fig. 1. The results on the right side present the IO curve with regression and estimated variability range (top left) as well as excitability estimations (bottom left) for every individual pulse in temporal order as in the MEP file and the corresponding estimated type-1 error (bottom right).

As indicated above, the different optimisation methods vary in their convergence and speed. The particle swarm will possibly take the longest computation time compared to simplex and interior-point methods but promises best convergence. The application cannot guarantee convergence for any of the methods. Therefore, users should graphically check the results. Furthermore, good convergence depends on the number of stimuli and their distribution throughout the entire curve. All methods need samples in the saturation regime, as well as for very weak stimuli to estimate the levels. The programme does not explicitly need several samples per stimulation strength as previously used in the literature. Furthermore, IO curves are known to show inter-pulse correlation and hysteresis effects, particularly for short inter-stimulus intervals [16]. Therefore, we recommend to fully randomise the stimulation strength during IO curve data acquisition.

The proper separation of the two variability parameters in the model with two variability sources, i.e., *v*_*x*_ and *v*_*y*_, requires more samples for a good result than the single-variability model. The model furthermore needs sufficient samples also in the upper and lower plateaus for estimating their full distribution information. Without sufficient samples across the entire range, the model can diverge to rather poor solutions and provide poor reliability of the parameters. The method outputs the log-likelihood as well as the Schwarz Bayesian information criterion (BIC) to reveal if the samples suffice, to indicate under- and over-fitting, and to compare which of the models may be the better fit given the available data. The lower the BIC metric of a model for specific data with specific data quantity, the better this model describes these data compared to a model with larger BIC value. Thus, a lower BIC value particularly suggests that the addition of parameters of model features are statistically justified for the available samples their distribution across the entire range of stimulation strengths and quantity. The BIC can serve as quality-of-fit metric and indicate the position in the variability–bias trade-off.

## Conclusion

We designed this IO curve analyser with the aim of providing a user-friendly environment for the analysis of brain stimulation IO curves, e.g., with TMS or suprathreshold electrical stimulation, in clinical and experimental neuroscience. The application can efficiently analyse measurements and translate the cloud of stimulus–response pairs with varying stimulus strengths into parameters of a regression curves, which can subsequently serve as biomarkers.

Most importantly, the tool wants to reduce the risk of spurious results and effects due to inappropriate statistical models or methods, such as previously highly popular linear regression of MEP IO curves. The latter ignores the strongly skewed MEP distribution and in consequence leads to unbalanced residuals, systematic bias, as well as unstable outlier-dependent model parameters. Furthermore, this method allows the extraction of quantitative variability information and also splits the variability into the stimulation-strength-dependent contributions, which act on the effective stimulation strength and excitability, from the independent contributions, such as peripheral calcium fluctuations as in short-term fatigue effects.

## Author contributions

KM and SMG envisioned and designed the overall project. SMG developed the core model and optimisation code, and KM modified the code and designed the graphical user interface and the overall structure of the application. SMG supervised this project. KM and SMG wrote the text.

## Supplementary material

Supplementary document to this article can be found online (https://github.com/BIOMAKE/Variability_MEP_IO_Curve_App).

